# Pancreatic Gα_s_ ablation disrupts tissue architecture and YAP signaling and unveils a compensatory regenerative response

**DOI:** 10.64898/2026.04.20.718494

**Authors:** Martina Rossotti, Juan I. Burgos, Dana J. Ramms, Agustín Romero, Valeria Burgui, Martín Zelicovich, Silvio A. Traba, Ana C. Heidenreich, J. Silvio Gutkind, Santiago A. Rodríguez-Seguí

## Abstract

Diabetes mellitus is characterized by chronic hyperglycemia and loss of pancreatic β-cell function and mass. Current therapies focus on β-cell protection and regeneration, led by GLP-1 receptor agonists. The G protein α-subunit (Gα_s_) acts as a key signaling node downstream of numerous GPCRs, integrating diverse signals that impact β-cell mass and function. Elucidating the integrative role of pancreatic Gα_s_ signaling is thus crucial for understanding β-cell biology. Our map of the pancreatic Gα_s_-coupled GPCR landscape reveals sophisticated, cell-type-specific networks, positioning Gα_s_ as a central hub for intra-pancreatic communication. Previous studies in mice with β-cell-specific or whole-pancreatic Gα_s_ deletion demonstrated reduced β-cell mass, impaired insulin secretion, and glucose intolerance. The stronger phenotype in the whole-pancreas model—marked by α-cell expansion and abnormal distribution—points to a crucial role for Gα_s_ in differential control of postnatal α- and β-cell proliferation. Here, we analyze the organ-wide consequences of Gα_s_ deletion using pancreas-specific Gα_s_ knockout mice (PGsKO). Consistent with prior findings, PGsKO mice exhibit reduced weight gain from four weeks and severe diabetes due to decreased β-cell mass and concomitant α-cell expansion. Furthermore, Gα_s_ loss induces profound architectural and functional defects in the exocrine pancreas, linked to YAP reactivation in acinar cells. Importantly, we observed attempted β-cell regeneration in PGsKO mice. Although insufficient to reverse diabetes, our results delineate the full pancreatic phenotype that may facilitate these regenerative efforts and suggest that strategically biasing GPCR signaling network away from Gα_s_ could be a viable strategy to promote β-cell regeneration from other pancreatic cell types.

**ARTICLE HIGHLIGHTS:**

- Gα_s_ is a central signaling hub that integrates diverse GPCR inputs across pancreatic cell types, yet its organ-wide role remained poorly defined.
- We addressed how pancreas-wide Gα_s_ deletion disrupts both endocrine and exocrine compartments, and whether regenerative programs are engaged.
- Gα_s_ loss caused severe diabetes through β-cell loss and α-cell expansion, induced profound exocrine defects with YAP reactivation, and triggered attempted β-cell regeneration from ducts and potentially other cell types.
- Our findings suggest that strategically biasing GPCR signaling away from Gα_s_ could promote regeneration from non-β-cell sources, offering new therapeutic avenues for diabetes.

## INTRODUCTION

Insulin-secreting pancreatic β-cells are central to glucose homeostasis, and their deficit or dysfunction leads to insulin deficiency, hyperglycemia, and the onset of diabetes (1). Consequently, a major therapeutic goal is to preserve, replace, or regenerate functional β-cells through approaches such as pharmacological agents, stem cell-derived β-cells, or cellular reprogramming (2–4). The development of Glucagon Like Peptide 1 Receptor (GLP-1R) agonists has been particularly revolutionary in this field (5). The GLP-1R is a G protein-coupled receptor (GPCR) that signals through both Gα_s_ and Gα_q_, and recent studies suggest that biasing this downstream signaling can fine-tune the beneficial effects of receptor agonists on β-cell function (6; 7). Beyond its direct insulinotropic action, GLP-1R agonist treatment can promote β-cell regeneration by stimulating β-cell proliferation and facilitating α-to-β cell transdifferentiation (8). Other GPCRs, such as the glucagon receptor (GCGR) which primarily signals through Gα_s_, have also been reported to modulate β-cell regeneration, either alone or in combination with GLP-1R agonists (9).

Critically, recent evidence emphasizes the importance of intra-pancreatic and intra-islet signaling crosstalk in modulating β-cell function and regeneration (10). Given that many GPCRs signaling through Gα_s_ (e.g., GLP1-R, GIPR, GCGR) are ubiquitously expressed across multiple pancreatic cell types, a localized ligand stimulus can simultaneously activate these receptors in a coordinated manner. Therefore, delineating the Gα_s_-dependence profile at the whole-pancreas level is expected to provide key insights into the global regenerative cues controlled by this critical signaling hub.

The role of Gα_s_ in β-cells has been studied using conditional knockout models. Mice with β-cell-specific deletion of Gα_s_ (βGsKO), generated by crossing *Gnas* floxed mice with *Ins1-Cre* or *Ins2-Cre* lines, develop severe hyperglycemia, glucose intolerance, and hypoinsulinemia (11; 12). These studies demonstrated that βGsKO mice have smaller islets, reduced β-cell mass, and impaired β-cell maturation. These defects stem from a lower β-cell proliferation rate during the first postnatal weeks, a critical period for establishing an adequate β-cell population.

The relevance of Gα_s_ at earlier developmental stages has been extended using mice with pancreatic-wide deletion from organ specification (PGsKO), generated with *Pdx1-Cre* (13). PGsKO mice become hyperglycemic by four weeks of age due to reduced β-cell mass and maturation defects. Unlike βGsKO mice, PGsKO islets exhibit a higher proportion of α-cells, suggesting a cell-type-specific role for Gα_s_ in controlling proliferation. It has been reported that PGsKO mice have a larger pancreas with ductal enlargements and a malabsorption phenotype, although the underlying mechanisms remain unexplored.

Building on this foundation, our study leverages the PGsKO model to provide a comprehensive analysis of the organ-wide consequences of Gα_s_ deletion. We first confirm the established diabetic phenotype resulting from Gα_s_ loss. To understand the signaling framework underlying these effects, we systematically mapped the Gα_s_-coupled GPCRs expressed across pancreatic cell types. We then characterize the associated islet cell abnormalities and extend these findings by uncovering profound defects in the exocrine compartment linked to YAP reactivation. Furthermore, we document attempts at β-cell regeneration within this altered pancreatic environment. Collectively, our work delineates the complete pathological landscape resulting from pancreatic Gα_s_ deficiency and suggests that strategically biasing GPCR signaling away from Gα_s_ could promote β-cell regeneration from other pancreatic cell types.

## RESEARCH DESIGN AND METHODS

### Mice

Mice with loxP sites flanking Gα_s_ exon 1 (*Gα*_s_ ^flox/flox^) (14) were maintained on a Black Swiss background. *Pdx1-Cre*^+/-^ mice (#014647) were backcrossed to C57BL/6 for ≥3 generations. Female *Gα*_s_ ^flox/flox^ mice were mated with male *Gα*_s_ ^flox/+^: *Pdx1-Cre*^+/-^ mice to generate pancreatic Gα_s_ knockout mice (PGsKO; *Gα*_s_ ^flox/flox^: *Pdx1-Cre*^+/-^). *Gα*_s_ ^flox/flox^ littermates lacking Cre were used as controls Genotyping was performed by PCR on tail DNA (primers in **Supplementary Table 1**) using DreamTaq polymerase under the following conditions: 95°C 4 min; 35 cycles (95/60-64/72°C 30 s each); 72°C 5 min. Mice were housed on a 12:12 h light/dark cycle with standard diet (2014S Teklad Global) and procedures approved by the University of Buenos Aires CICUAL (protocol 144B/2024), following ARRIVE guidelines. Experimental procedures and tissue collection were performed at indicated time points, with birth considered postnatal day 0. Animals were euthanized by cervical dislocation.

### Intraperitoneal Glucose Tolerance Test (IPGTT)

Blood samples for glucose detection were collected from the tail vein, and measured using a hand-held OneTouch Ultra glucometer (LifeScan, Milpitas, CA, USA). To perform IPGTT, animals were fasted for either 6 daily hours or overnight prior to the glucose tolerance test. Fasting duration was not considered an independent experimental variable. Nevertheless, its potential impact on baseline glycemia and overall glucose response was evaluated, and no significant differences were observed between fasting regimens. Therefore, data were analyzed together. A glucose solution was administered by intraperitoneal injection at 2 mg/g, and blood glucose levels were measured at 15, 30, 60 and 120 min after the glucose loading. If the glucose level was higher than 600 mg/dl (the glucometer’s upper limit), the value of 600 mg/dl was recorded.

### Immunofluorescence and Morphometric Analysis

Pancreases from 10-week-old mice were fixed in 4% paraformaldehyde overnight at 4°C, paraffin-embedded, sectioned at 8 μm, and stained following previously reported immunodetection protocols (3; 15). Briefly, after rehydration, heat-mediated antigen retrieval was performed in 0.01 M sodium citrate pH 6.0 for 5 min. Sections were permeabilized with 0.5% Triton X-100 for 30 min, blocked with 3% donkey serum, and incubated overnight at 4°C with primary antibodies: rat anti-E-cadherin (Invitrogen 13-1900), guinea pig anti-insulin (DAKO IS002 or Linco 4011-01), mouse anti-glucagon (Sigma G2654), rabbit anti-amylase (Sigma A8273), rabbit anti-laminin (Sigma L9393), hamster anti-Mucin1 (Thermo Fisher MA5-11202), and guinea pig anti-Pdx1 (Abcam ab47308). Slides were washed and incubated for 45 min at 4°C with DBA-Fluorescein (Vector Laboratories FL-1031-2) or appropriate secondary antibodies, followed by DAPI (Thermo Fisher 62248) for 15 min. Images were captured on a Zeiss LSM 900 confocal microscope.

For morphometry, 2-3 sections (>100 μm apart) per animal were analyzed using ImageJ. β-cell fraction was calculated as (insulin-positive area/total pancreatic area) × 100. Islet size and acinar cell area were measured in calibrated units. For acinar cell area quantification, cells within a 20-cell radius of islets were compared to distant cells. For E-cadherin quantification, confocal images were acquired with identical parameters. Islet and adjacent acinar ROIs were manually delineated, and mean fluorescence intensity was measured after uniform background subtraction, with islet signal normalized to adjacent acinar tissue. For cell counts, 3-5 equally spaced sections covering the entire pancreas were imaged, and positive cells were counted manually from ≥3 mice per group.

### H&E Staining

Paraformaldehyde-fixed (4%), paraffin-embedded pancreatic sections (5 μm) were stained with hematoxylin and eosin. Whole-slide images were acquired on an AT2 Aperio ScanScope and analyzed with QuPath software (version 0.2.3).

### Compilation of Gαs-coupled GPCRs

GPCRs with documented Gαs coupling were compiled from the GPCRdb database (raw data file *GPCR-G_protein_couplings.xlsx* accessed 1 December 2025 from https://github.com/protwis/gpcrdb_data). Candidate receptors were identified by merging entries from the “GtoPdb Gs family,” “Bouvier Gs family,” and “Marteinsson-Myanov Gs family” datasets. The list was expanded with receptors annotated for primary Gαs coupling in the “Lambert-RGB-GDP_Econst” table and validated against the GtoPdb “Primary Transducer” field, yielding 78 unique mouse GPCR genes (**Supplementary Table 2**).

### Single-Cell RNA-Seq Analysis

We reanalyzed an integrated dataset of six public mouse pancreatic islet scRNA-seq studies (16), normalized and clustered using Conos. GPCR expression was assessed in β, α, ductal, and acinar clusters. Genes expressed in ≥1% of cells within any cluster (n=23) were visualized by dot plot, hierarchically ordered favoring β-cell expression, with color indicating mean log-normalized expression and dot size indicating the percentage of expressing cells.

### Bulk RNA-seq and ChIP-seq Data Processing

Raw sequencing data (**Supplementary Table 3**) were obtained from the SRA. RNA-seq reads were aligned to the mouse genome (mm39) using HISAT2 v2.2.1 (17); ChIP-seq reads were aligned with Bowtie2 v2.5.1 (18). SAM files were converted to BAM and sorted using SAMtools (19). For visualization in IGV (20), normalized coverage tracks were generated: for RNA-seq, using deepTools (21) bamCoverage (bin size 1 bp, CPM normalization); for ChIP-seq, using igvtools count (-w 25 -e 250 -f mean -z 7).

### Quantification and Statistical Analysis

Data normality was assessed with the Shapiro-Wilk test. Gaussian data are presented as mean ± SEM and analyzed by unpaired two-tailed Student’s t-test (two groups), or one-way or two-way ANOVA with Dunnett T3, Tukey, or Bonferroni post-hoc tests (≥3 groups). Non-Gaussian data are presented as median (interquartile range) and analyzed by Mann-Whitney (two groups) or Kruskal-Wallis with Dunn’s post-hoc test (≥3 groups). Multinomial regression with Benjamini-Hochberg correction was used for cell proportion analyses. p<0.05 was considered significant. Analyses were performed using GraphPad Prism 9.0 or R. Specific statistical tests for each figure are provided in the corresponding figure legends.

### Data and Resource Availability

Supporting data are available from the corresponding author on reasonable request.

## RESULTS

### PGsKO Mice Display Impaired Weight Gain and Severe Diabetes From Four Weeks of Age

To assess the global effects of pancreatic Gα_s_ loss, we generated PGsKO mice (Gα_s_^fl/fl^: Pdx1-cre^+^) by mating Gα_s_^fl/fl^ females with Gα_s_^fl/+^:Pdx1-Cre^+/-^ males (**Fig. 1*A***; **Supplementary Fig. 1*A***). Littermate Gα_s_^fl/fl^ mice were used as controls. PGsKO mice exhibited severe postnatal growth retardation, apparent from four weeks of age, leading to a significantly lower body weight than controls (**Fig. 1*B*** and ***C***). Despite this reduced body weight, PGsKO mice consistently displayed an enlarged pancreas (**Supplementary Fig. 1*B***), a feature that has also been reported in earlier studies of Gα_s_ deletion. Consistent with previous observations, PGsKO mice were hyperglycemic by four weeks of age (**Fig. 1*D*** and ***E***), and the diabetic phenotype progressed to severe hyperglycemia by ten weeks (**Fig. 1*F*** and ***G***).

**Figure 1.**
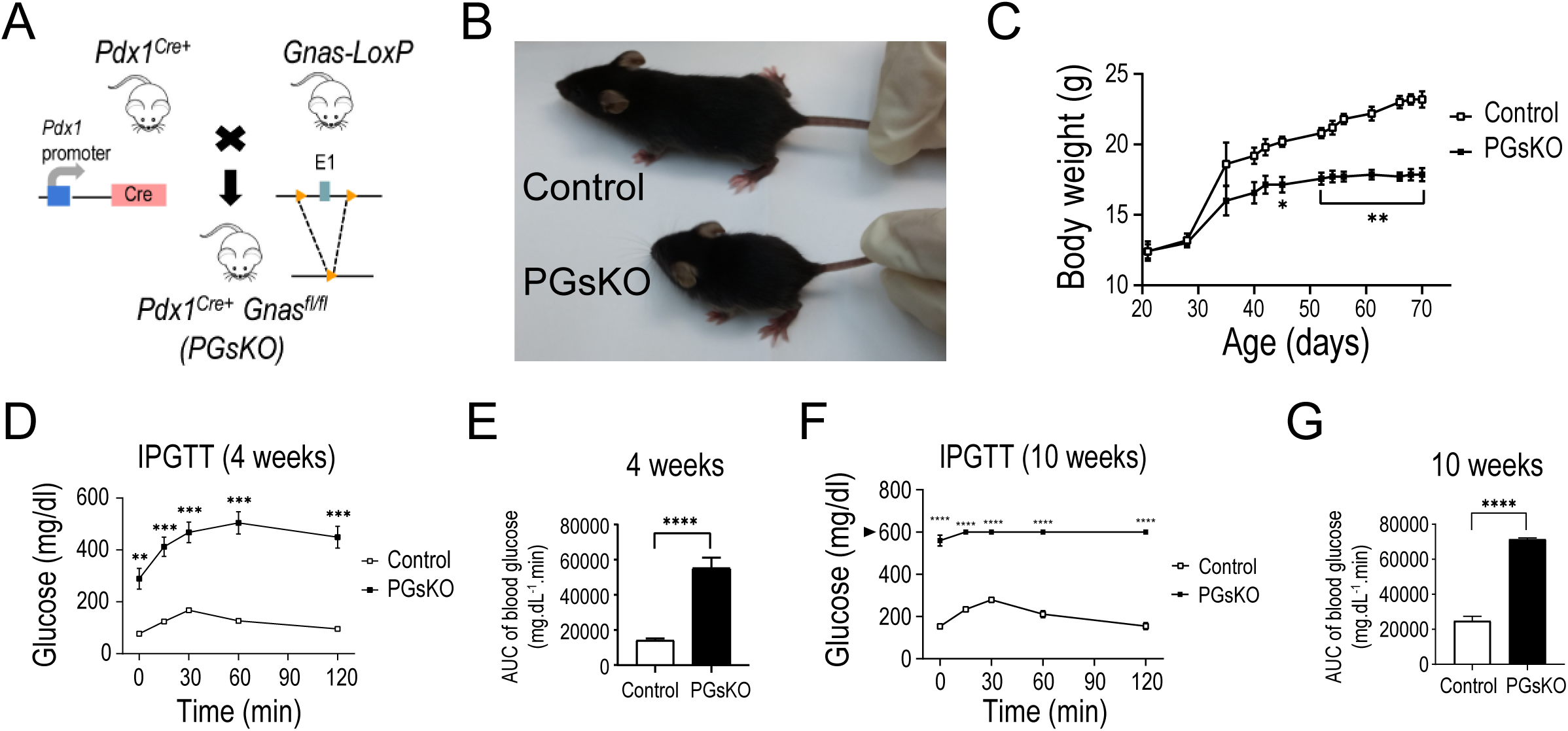
PGsKO mice display impaired weight gain and severe diabetes from four weeks of age. ***A***: Experimental strategy to generate pancreas-specific *Gnas*-knockout mice, hereafter PGsKO. ***B*:** Photograph of 5-week-old control and PGsKO mice. ***C*:** Growth curves of control and PGsKO mice (*n* = 6 mice/group). ***D, F*:** Plasma glucose levels during the intraperitoneal glucose tolerance test in control and PGsKO mice at 4 and 10 weeks of age (IPGTT, 2mg/g; arrowhead indicates the maximum limit detected by the glucometer). Panel ***E*** shows the area under each curve (AUC) for 4 weeks of age and Panel ***G*** for 10 weeks of age control and PGsKO mice (*n* = 7 mice/group). Data are expressed as mean ± SEM. Statistical analysis was conducted by multiple t-test with Bonferroni correction, and two-tailed Welch’s t-tests were applied for AUC graphs. *p < 0.05, **p < 0.01, ***p < 0.001, ****p < 0.0001.

### The Gα_s_-coupled GPCR Landscape of the Mouse Pancreas

To define the signaling potential of the Gα_s_ pathway, we profiled the expression of GPCRs annotated for canonical or alternative Gα_s_ coupling across major pancreatic cell types. We reanalyzed integrated data from six public mouse pancreatic single-cell RNA-sequencing datasets (16) (**Fig. 2*A***) and cross-referenced findings with the GPCRdb database (22).

**Figure 2.**
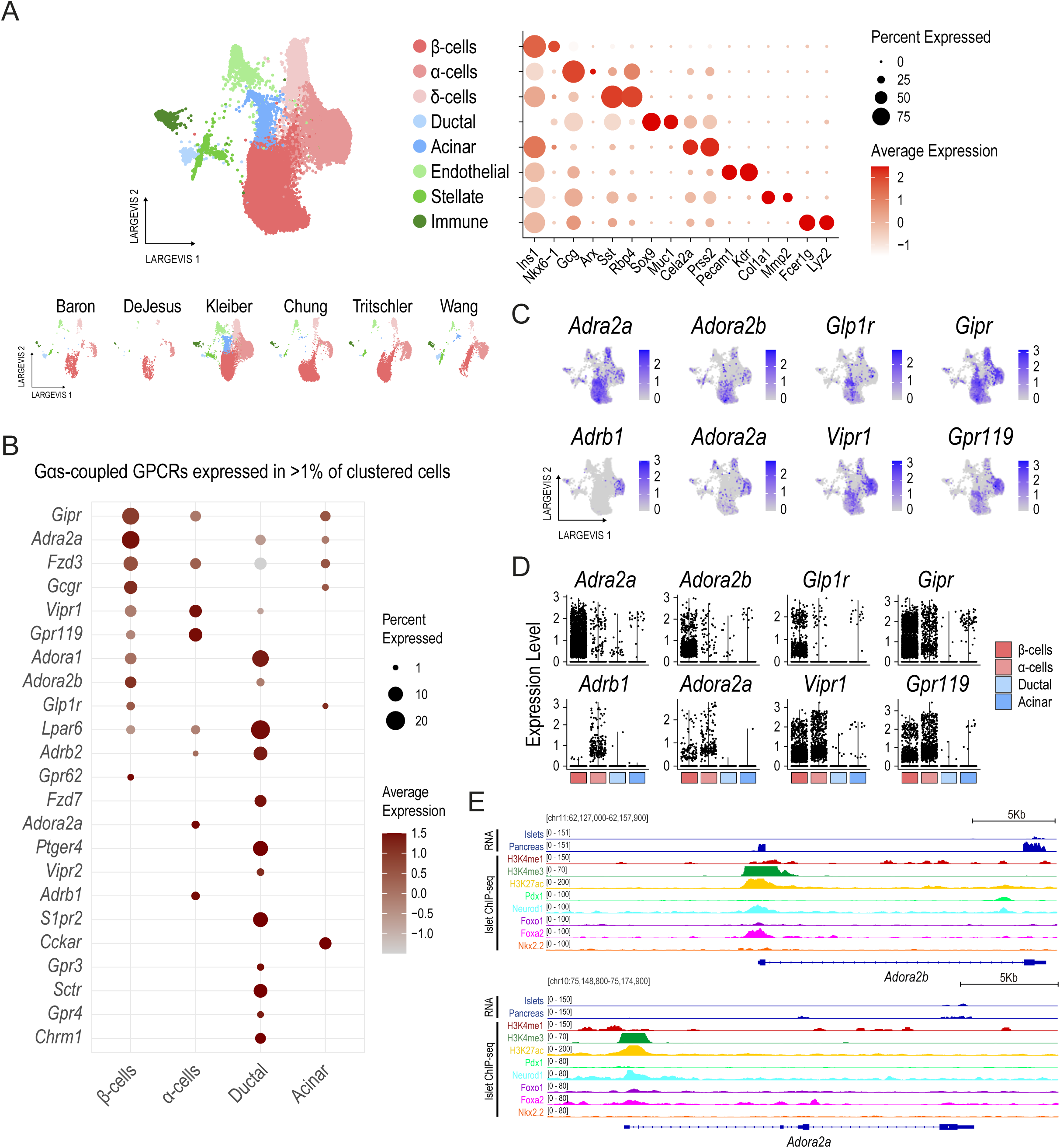
The Gα_s_-coupled GPCR landscape of the mouse pancreas. ***A***: UMAP (left panels) and dot (right panel) plots of single-cell transcriptomes profiled from six different mouse adult pancreatic islet datasets, integrated with Conos. UMAP plots at the bottom show cell distribution according to the sample of origin. The dot plot shows the expression of key markers for endocrine, ductal, acinar, stellate, immune, and endothelial cell types, used to define clusters. Colour intensity indicates mean normalized expression, and dot size indicates the percentage of expressing cells. ***B*:** Dot plot showing the expression of Gα_s_-coupled GPCRs that are expressed in >1% of α-, β-, ductal or acinar clustered cells. Genes are ordered by a hierarchical score favoring β-cell expression. ***C, D*:** Feature (*D*) and Violin (*E*) plots showing the expression of selected Gα_s_-coupled GPCRs. ***E*:** Gene expression and chromatin features at *Adora2b* and *Adora2a* loci. Bulk RNA-seq data confirm enriched expression of both *Adora2b* and *Adora2a* in islets compared to whole pancreas. Chromatin profiling of islets (transcription factor binding, H3K4me3, H3K4me1, H3K27ac) reveals active promoter and enhancer architecture at each locus, supporting their endocrine-specific transcriptional regulation. For all expression plots (*B*, *D*, *E*), expression levels are shown in log-normalized counts.

Our analysis revealed a rich landscape of Gα_s_-coupled GPCRs with distinct, cell-type-specific expression patterns. Of 78 GPCRs annotated for primary or alternative Gα_s_ coupling (**Supplementary Table 2**), we detected significant expression (in >1% of cells per cluster) for 23 receptors (**Fig. 2*B***). As expected, β-cells prominently expressed key metabolic regulators such as *Glp1r*, *Gipr*, and *Gcgr*. Ductal cells exhibited the largest proportion of selectively enriched receptors, including *Ptger4*, *S1pr2*, *Sctr* and *Chrm1*. The *Cckar* receptor was identified as specific to acinar cells.

Beyond cataloguing expression, our map revealed sophisticated signaling architectures that suggest precise physiological control, particularly within islet cells. We identified a mirrored receptor distribution for two neuromodulatory axes, each employing a distinct logic to fine-tune endocrine output.

The adrenergic system employs opposing G-protein couplings to orchestrate divergent hormonal responses. In β-cells, the inhibitory receptor *Adra2a* (canonically Gα_i_) was enriched, whereas α-cells selectively expressed the stimulatory Gα_s_-coupled receptor *Adrb1* (**Fig. 2*C*** and ***D***). Notably, while *Adra2a* is annotated for primary Gα_i/o_ coupling, it can also signal through Gα_s_ under certain contexts (22), suggesting that catecholamine effects in β-cells may involve complex, context-dependent modulation rather than pure inhibition. This arrangement supports a model in which sympathetic catecholamines suppress insulin secretion while promoting glucagon release, thereby raising blood glucose during stress or energy demand (23).

In contrast, the adenosine system relies on parallel Gα_s_-coupled receptors to modulate endocrine output. The Gα_s_-linked receptors *Adora2a* and *Adora2b*, both capable of elevating intracellular cAMP (24; 25), were partitioned to α- and β-cells, respectively (**Fig. 2*C*** and ***D***). Analysis of the pancreatic islet epigenomic landscape (**Fig. 2*E***) confirmed a tissue-specific regulatory architecture in α- and β-cells, with active chromatin at these loci marked by distinct promoter and enhancer signatures co-bound by islet-specific transcription factors. This organization suggests a framework in which local adenosine signals can concurrently enhance glucagon release and modulate insulin secretion through receptor-specific, context-dependent mechanisms.

This pattern reveals a fundamental design principle: Gα_s_ mediates conflicting signals in β-cells but convergent signals in α-cells. In β-cells, Gα_s_-coupled receptors (*Glp1r*, *Adora2b*) must elevate cAMP to counterbalance the potent inhibitory drive imposed by the canonically Gα_i_-coupled receptor *Adra2a*. In α-cells, by contrast, Gα_s_-coupled receptors (*Adrb1*, *Adora2a*) converge to amplify stimulatory signaling without an opposing Gα_i_-dominated input among the receptors profiled.

We also identified receptors with shared endocrine roles, such as *Vipr1* and *Gpr119* expressed in both α- and β-cells. This comprehensive map positions Gα_s_ as a critical node for integrating diverse hormonal, paracrine, and neuronal inputs within the pancreatic epithelium.

Collectively, this GPCR landscape provides a molecular blueprint for the severe pancreatic dysfunction observed upon Gα_s_ deletion. The ablation of Gα_s_ in PGsKO mice is predicted to cripple the essential stimulatory arms in β-cells, leading to insulin deficiency, while simultaneously disrupting the convergent stimulatory tuning in α-cells, promoting dysregulated glucagon output—a combination that directly explains the severe hyperglycemia. Having defined this logic-driven endocrine signaling network, we next sought to determine the functional consequences of ablating its central integrator and to investigate whether the loss of Gα_s_ in ductal and acinar compartments contributes to the diabetic phenotype through broader tissue crosstalk.

### Loss of Gα_s_ Leads to Reduced β-Cell Mass and Expansion of the α-Cell Compartment

It was previously reported that PGsKO mice have smaller islets with reduced β-cell mass and increased α-cell numbers, based on initial immunohistochemical analysis (13). To build on these qualitative observations, we performed a detailed quantification of these phenotypes in the pancreata of 10-week-old PGsKO and control mice. Immunostaining for insulin and glucagon confirmed a significant disruption of islet architecture in PGsKO mice, characterized by α-cells occupying the core of the islet—a stark contrast to their normal peripheral localization (**Fig. 3*A***).

**Figure 3.**
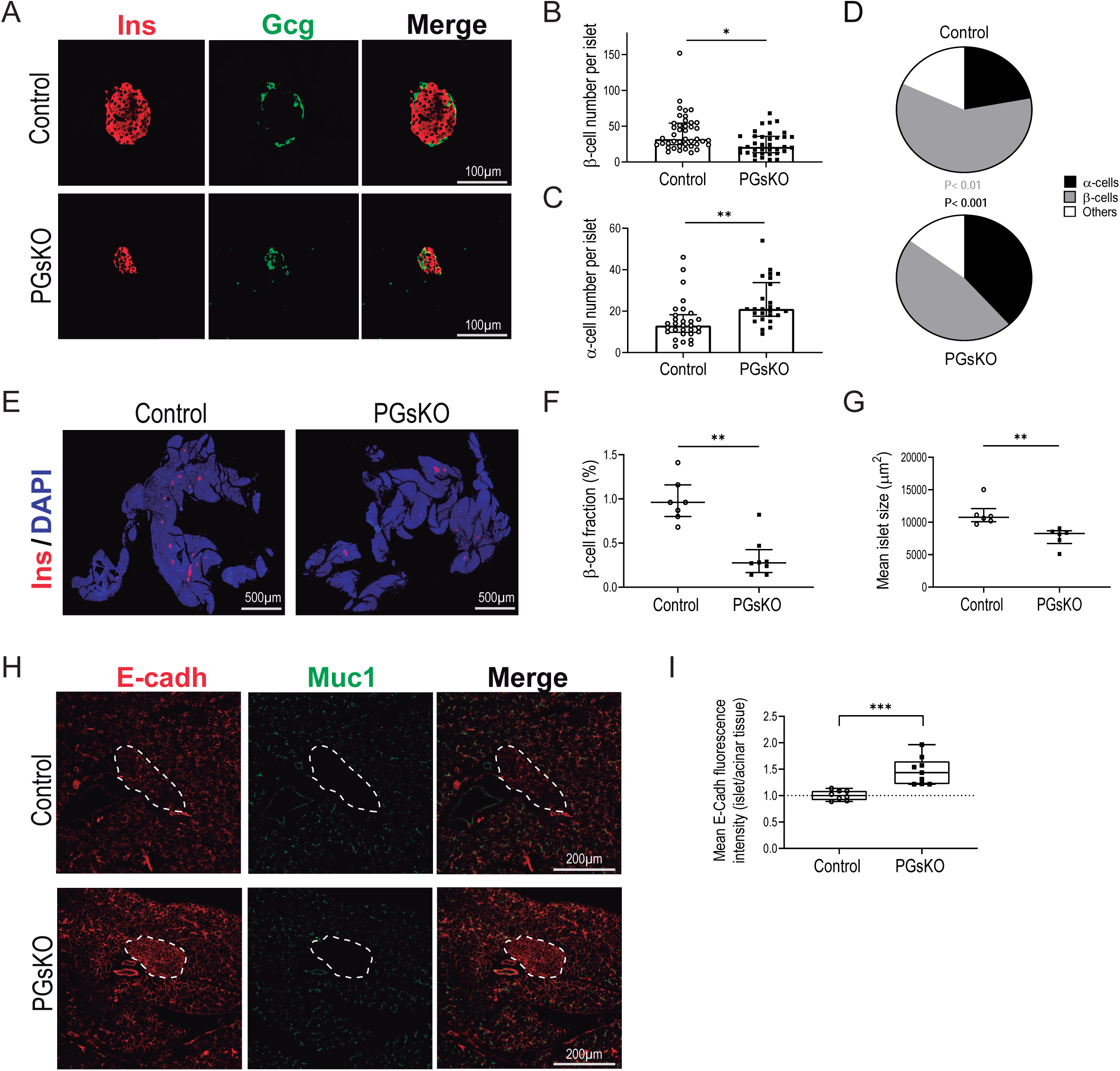
Loss of Gα_s_ leads to reduced β-cell mass and expansion of the α-cell compartment. ***A***: Representative image of an islet immunostained for insulin and glucagon in 10-week-old PGsKO and control mice. ***B-D*:** Quantification of the number of β-cells (*B*) and α-cells (*C*) per islet, and the proportion of α, β, and other endocrine cell types in the pancreatic islets of PGsKO and control mice (*D*) (*n* = 4 mice/group). ***E*:** Representative images of insulin and DAPI immunostaining of whole pancreatic sections of 10-week-old PGsKO and control mice. ***F:*** Pancreatic β-cell fraction (*n*= 7 control and *n*=8 PGsKO mice) and (**G**) mean islet size of 6 pancreatic slides from each animal (*n*= 6 mice/group). ***H*:** Fluorescent staining of pancreatic sections from PGsKO and control mice (10 weeks old), highlighting E-cadherin and mucin-1 expression. White dashed lines outline the islets. ***I*:** Mean E-cadherin fluorescence intensity in islets normalized to acinar tissue levels in 10-week-old PGsKO and control mice, presented as mean with bars indicating minimum and maximum values. Statistical significance was assessed using a two-tailed Welch’s t-test (*n*= 8 control and 9 PGsKO mice). Data are expressed as the median (interquartile range) in (*B*), (*C*), (*F*), and (*G*) where statistical analysis was conducted by the Mann-Whitney test (two-tailed). A multinomial regression comparison for each cell fraction with Benjamini-Hochberg correction was performed in (*D*). *p < 0.05, **p < 0.01, ***p < 0.001.

Quantitative analysis revealed that PGsKO islets contained, on average, 33% fewer β-cells (**Fig. 3*B*** and ***D***) and 33% more α-cells than controls (**Fig. 3*C*** and ***D***). Strikingly, while the average β-cell count per islet was two-thirds that of controls, the global β-cell fraction across the entire pancreas was reduced by 75% (**Fig. 3*E*** and ***F***). This disproportionate loss is explained by a combination of fewer total islets and a 25% reduction in average islet size in PGsKO mice (**Fig. 3*G***).

Furthermore, immunofluorescence for the cell adhesion marker E-Cadherin and the polarity marker Mucin1 revealed altered islet organization. Islets from PGsKO mice exhibited intensified E-Cadherin staining, suggesting stronger intra-islet cell adhesion interactions (**Fig. 3*H*** and ***I***; **Supplementary Fig. 2**).

### PGsKO Mice Exhibit Exocrine Tissue Architectural Defects, Aberrant Cell Polarization, and YAP Reactivation

A comprehensive analysis of pancreatic Gα_s_ loss revealed profound architectural and functional defects in the exocrine compartment. Histological examination of H&E-stained sections indicated that Gα_s_ deletion induces acinar cell hypertrophy and a loss of cellular polarity. This was evidenced by the aberrant cytoplasmic distribution of eosinophilic zymogen granules, which were dispersed throughout the cell in PGsKO mice, in contrast to their strict apical localization in controls (**Fig. 4*A***).

**Figure 4.**
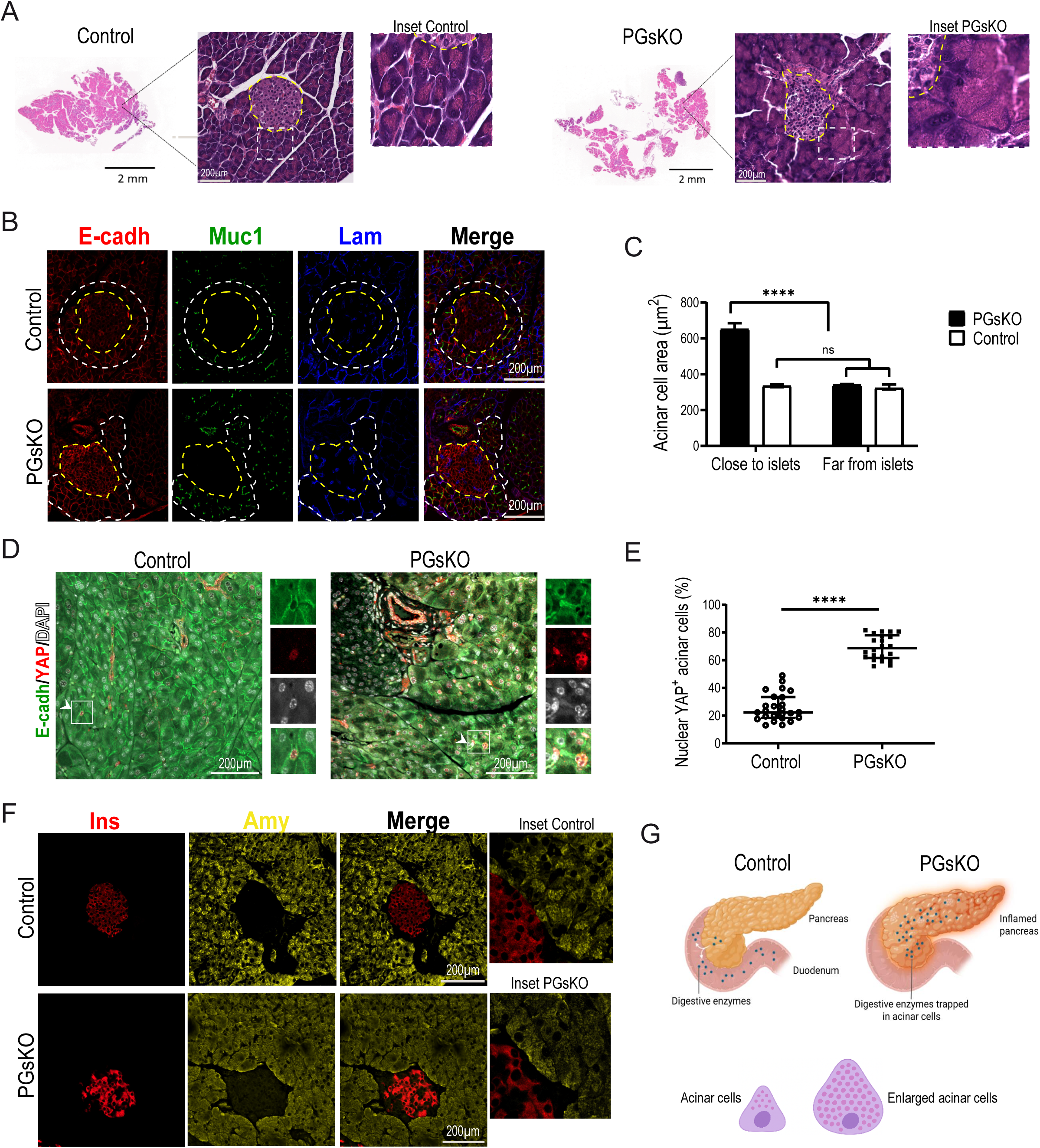
PGsKO mice show exocrine tissue architectural defects, with abnormal cell polarization and YAP reactivation. ***A***: Representative Hematoxylin and Eosin-stained sections of whole adult pancreas from control and PGsKO mice, with their corresponding magnified views. Yellow dashed lines outline the islets, while white dashed boxes indicate acinar cells immediately adjacent to the islets. ***B*:** Fluorescent labeling of E-cadherin, mucin-1, and laminin in pancreatic tissue from 10-week-old PGsKO and control mice. Yellow dashed lines outline the islets, while white dashed lines indicate acinar cells immediately adjacent to the islets. ***C*:** Quantification of acinar cell area directly adjacent (within 20-cell radius) or distant from the islets in pancreatic sections of PGsKO and control mice (*n* = 3 mice/group). **D:** Fluorescent staining of pancreatic sections from PGsKO and control mice (10-weeks-old), labeling E-cadherin, YAP and DAPI; white arrowheads and insets highlight nuclear YAP expression. ***E:*** Quantification of the percentage of acinar cells with nuclear YAP positive staining per pancreatic section in PGsKO and control mice (*n* = 4 mice/group). ***F*:** Representative images of insulin and amylase immunostaining of pancreatic slides of 10-week-old PGsKO and control mice, with their corresponding magnified views. ***G:*** Schematic representation of the proposed alterations in the exocrine pancreas of PGsKO mice and their corresponding acinar cell phenotypes in the absence of Gα_s_ compared to control mice. Data are presented as median (interquartile range) and were analyzed by Kruskal-Wallis test, followed by the Dunn multiple comparisons test among three or more groups in ***C*** and Mann-Whitney test (two-tailed) in ***E***. ****p < 0.0001, ns= not significant.

Immunostaining for cell polarity markers—E-Cadherin (lateral adhesion), Mucin 1 (apical polarity), and Laminin (basal membrane)—further confirmed this disrupted organization. Unexpectedly, quantification of acinar cell size revealed that cells in close proximity to islets were larger than those located further away (**Fig. 4*B*** and ***C***). This spatially localized defect suggests a disruption of normal islet–acinar paracrine communication, consistent with impaired Gα_s_-mediated signaling networks mapped in **Fig. 2**.

We next investigated a potential mechanism for this phenotype. Given that epidermal-specific loss of Gα_s_ activates YAP and drives tumorigenesis in skin (26), we hypothesized a similar reactivation could occur in the pancreas. Although YAP plays a key role in progenitor cells during pancreas development (15), its activity in the adult organ is highly restricted, with significant nuclear localization largely confined to ductal and centroacinar cells (27), while being suppressed in mature acinar and endocrine cells. We therefore evaluated its status in the PGsKO model.

Strikingly, while YAP expression in control pancreases was restricted to ductal and centroacinar cells (**Fig. 4*D***, Control, white arrowheads and inset; **Supplementary Fig. 3*A***), it was robustly reactivated in the hypertrophic acinar cells of PGsKO mice, with prominent nuclear localization (**Fig. 4*D***, PGsKO, white arrowheads and inset; **Supplementary Fig. 3*A***). On average, the number of acinar cells with nuclear YAP in PGsKO pancreases was 2.6-fold higher than in controls (**Fig. 4*E***).

Further supporting our H&E observations and indicating acinar cell malfunction, immunofluorescence revealed an aberrant cytoplasmic distribution of amylase in PGsKO mice. This presented as a diffuse, lower-intensity signal compared to the strong, apically localized pattern in controls (**Fig. 4*F***; **Supplementary Fig. 3*B***). These results are consistent with aberrant zymogen granule retention within dysfunctional, enlarged acinar cells (**Fig. 4*G***).

### β-Cell Regeneration is Increased in the Pancreas of PGsKO Mice

Our initial characterization established that PGsKO mice have smaller islets (**Fig. 3*G***). A detailed quantification of islet size distribution revealed that this difference was primarily due to a significant increase in the proportion of very small islets, consisting of only 4-13 cells (**Fig. 5*A***). We also observed a greater number of isolated β-cells, as well as very small cell clusters (<4 cells), dispersed within the acinar tissue; however, the majority of these extra-islet insulin-producing cells (IPCs) were situated adjacent to, or immediately surrounding, ducts (**Fig. 5*B*** and ***C***; **Supplementary Fig. 3*B***).

**Figure 5.**
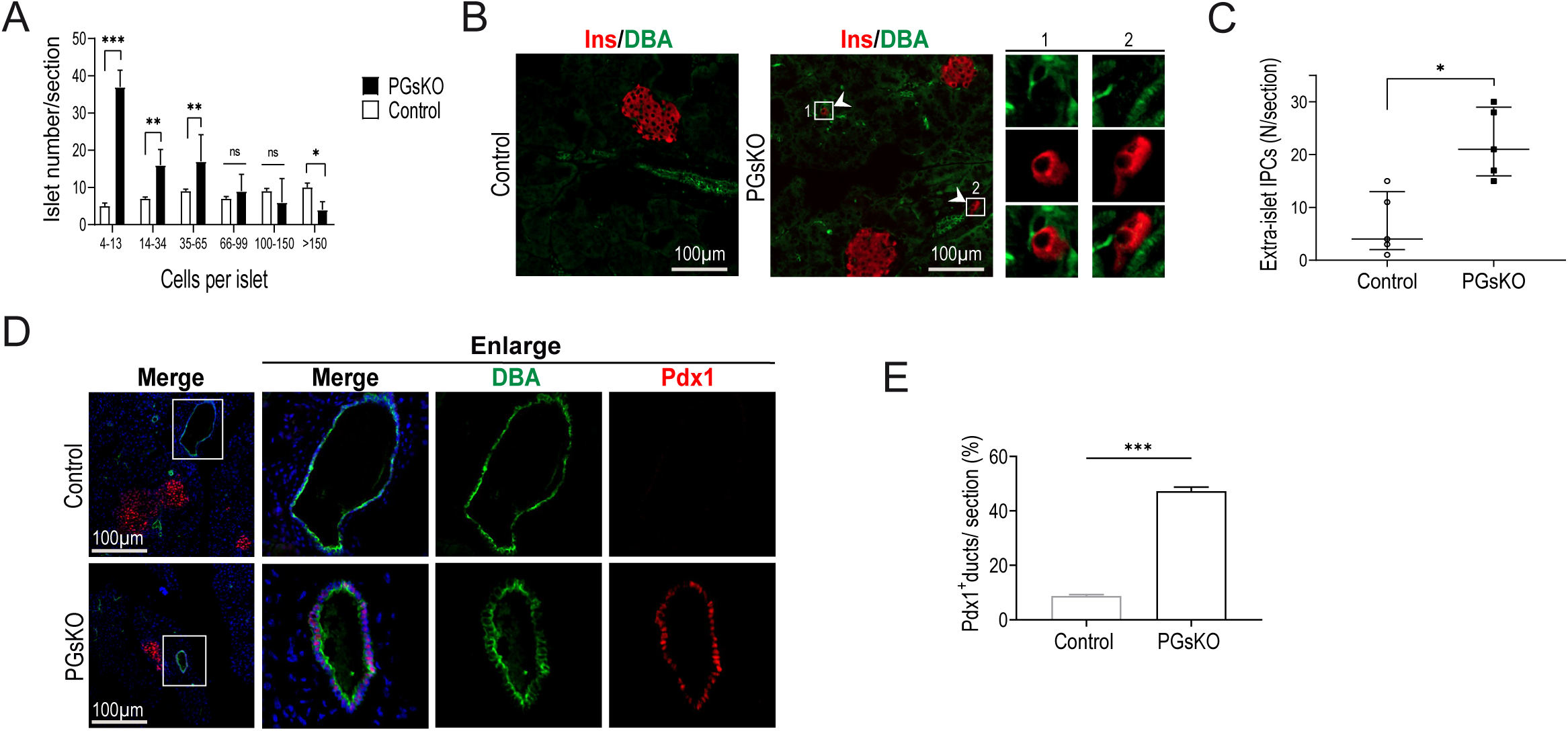
β-cell regeneration is increased in the pancreas of PGsKO mice. ***A***: Islet size distribution in pancreatic sections of 10-week-old PGsKO and control mice (*n* = 4 mice/group). ***B:*** Representative insulin and dolichos biflorus agglutinin (DBA) immunostaining in pancreatic sections from 10-week-old control and PGsKO mice. White arrowheads and insets (PGsKO) highlight extra-islet insulin-producing cells (IPCs). ***C:*** Quantification of extra-islet IPCs per pancreatic section of PGsKO and control mice (*n* = 5 mice/group). ***D:*** Fluorescent staining of pancreatic sections from PGsKO and control mice (10-week-old), labeling Pdx1, DBA and DAPI with their corresponding magnified views of ducts. ***E:*** Quantification of the percentage of ducts with nuclear Pdx1 positive staining per pancreatic section in PGsKO and control mice (*n* = 3 mice/group). In ***A***, data are expressed as mean ± SD and statistical analysis was conducted by using multinomial regression comparison of each category with Benjamini-Hochberg correction for multiple tests. In ***C*** data are expressed as median (interquartile range) and were analyzed by the Mann-Whitney test (two-tailed), and in ***E***, data are expressed as mean ± SEM where statistical analysis was conducted by two-tailed Welch’s t-tests. *p < 0.05, **p < 0.01, ***p < 0.001, ns= not significant.

Despite the severe diabetic state of PGsKO mice, the presence of these nascent endocrine cell clusters is consistent with attempted β-cell regeneration. This may occur through the transdifferentiation of other pancreatic cell types or the proliferation of pre-existing β-cell precursors (4).

Further supporting a potential contribution from the ductal compartment to β-cell neogenesis in PGsKO mice, immunostaining revealed widespread re-expression of the key developmental transcription factor Pdx1 in ductal cells (**Fig. 5*D*** and ***E***; **Supplementary Fig. 4**), suggesting reactivation of their endocrine-fate competence.

## DISCUSSION

This study provides a comprehensive analysis of pancreatic Gα_s_ deletion, revealing its essential role as a master regulator of organ-wide homeostasis in the adult mouse. We confirm that PGsKO mice develop severe growth retardation and progressive diabetes, and we establish that Gα_s_ loss triggers a triad of profound defects: disrupted cellular architecture, ectopic YAP reactivation, and a compensatory—albeit insufficient—regenerative response.

While mice with β-cell-specific Gα_s_ deletion (βGsKO) are hyperglycemic (11; 12), the PGsKO model presents a stronger and more complex phenotype. The onset of severe growth retardation and hyperglycemia at four weeks, which worsens by ten weeks, suggests that the critical function of Gα_s_ in maintaining metabolic homeostasis occurs postnatally. Although we cannot rule out earlier roles in pancreatic progenitors, the postnatal emergence of the phenotype indicates that either developmental compensation occurs or that Gα_s_ is critical for maintaining—rather than establishing—pancreatic integrity. This is consistent with its function as a central node in the extensive postnatal GPCR signaling network we have mapped.

A central finding of our work is the organ-wide dysregulation stemming from Gα_s_ loss. The exocrine compartment exhibited acinar cell hypertrophy, a complete loss of polarity evidenced by mislocalized zymogen granules and amylase, and the reactivation of the Hippo pathway effector YAP—a known driver of proliferation and hypertrophy. This YAP reactivation, previously documented in Gα_s_-deficient skin stem cells (26), now emerges as a conserved mechanism in the pancreas, directly linking Gα_s_ signaling to the control of acinar cell size and identity. However, while our results revealed an enlarged acinar compartment and ductal abnormalities that drive profound architectural tissue defects, we did not observe malignant transformation in PGsKO mice, in contrast to the reported tumorigenesis following Gα_s_ deletion in epidermal cells (26). Interestingly, pancreatic carcinogenesis has been associated with constitutive Gα_s_ activation in a KRAS-driven context (28), underscoring the critical, tissue-specific outcomes of Gα_s_ pathway modulation. Notably, acinar hypertrophy in PGsKO mice was most pronounced near islets, suggesting that the loss of Gα_s_ disrupts normal paracrine crosstalk, a concept strongly supported by our GPCR map, which shows numerous shared and cell-type-specific receptors.

The architectural defects extended to the islets, which displayed significantly altered adhesion properties and cell type composition. We observed a pronounced increase in E-cadherin levels within PGsKO islets. This suggests a potential strengthening of intra-islet cell adhesion, a phenomenon that may serve as a compensatory mechanism to preserve islet integrity amidst widespread tissue dysfunction. This finding aligns with studies showing that E-cadherin is essential for maintaining exocrine tissue architecture postnatally (29) and that coordination between ECM and cell-cell adhesion governs islet aggregation and maturation (30). The altered polarity in acinar cells, likely stemming from disrupted adhesion, thus appears to have ripple effects on the endocrine compartment, underscoring the interdependence of the entire organ.

Our GPCR expression analysis provides a mechanistic framework for the severe endocrine dysfunction. The transcriptomic landscape reveals a fundamental design principle: Gα_s_ mediates conflicting signals in β-cells (where stimulatory receptors like GLP1R and ADORA2B must counteract inhibitory ADRA2A) but convergent signals in α-cells (where ADRB1 and ADORA2A synergize to amplify secretion). The ablation of Gα_s_ therefore cripples essential stimulatory inputs in β-cells, causing insulin deficiency, while simultaneously disrupting the precise cAMP-dependent tuning in α-cells, promoting dysregulated glucagon output. Together, these defects provide a coherent explanation for the severe hyperglycemia observed in PGsKO mice.

The complexity of this signaling disruption may be further amplified by the inherent promiscuity of key GPCRs. Database annotations indicate that *Adra2a*, although canonically Gα_i_-coupled, can display context-dependent Gα_s_ coupling, while the canonically Gα_s_-coupled receptors *Adora2a*, *Adora2b*, *Adrb1* and *Glp1r* can engage Gα_q/11_ or Gα_i/o_ pathways (22). In the PGsKO model, the absence of Gα_s_ could therefore not only delete specific pathways but also “skew” the output of these promiscuous receptors toward their remaining alternative G proteins, potentially exacerbating the hormonal imbalance.

Critically, the PGsKO phenotype stems not only from cell-autonomous loss of Gα_s_ but from the collapse of an intricate local signaling network. Our map includes receptors for numerous locally-produced mediators—such as glucagon (GCGR) from α-cells (31), cholecystokinin (CCKAR) whose ligand can be delivered from intrapancreatic neurons or the gut (32), and prostaglandins (PTGER2/4) and adenosine (ADORA receptors) generated within the pathological pancreatic tissue by acinar or immune cells (33; 34). Gα_s_ deletion in all pancreatic epithelial cells therefore severs communication in both directions: it prevents acinar and ductal cells from receiving endocrine and neural signals via Gα_s_-coupled GPCRs (e.g., GLP-1 via GLP1R) and equally prevents islet cells from integrating paracrine cues from the exocrine compartment or neural/gut-derived inputs. This failure of intra-pancreatic and inter-organ crosstalk may act as a fundamental driver of organ-wide dysfunction.

This signaling collapse also provides a framework for understanding the expanded α-cell compartment and the signs of attempted β-cell regeneration. The combination of exocrine dysfunction and YAP activation may create a proliferative, regeneration-biased tissue environment that supports shifts in cell state. In addition, the profound β-cell loss and associated hyperglycemia could engage compensatory regenerative programs, including those involving GLP-1R–dependent α-to-β cell conversion (8). The presence of isolated β-cells and small endocrine clusters—particularly adjacent to ducts showing Pdx1 re-expression—is consistent with ongoing neogenic activity. This aligns with reports that β-cell formation from ductal epithelium can be induced in adult mice under certain stress or injury conditions (35). However, the failure of these processes to restore normoglycemia is likely rooted in the global loss of Gα_s_. Adenosine signaling is a potent inducer of β-cell proliferation (24), and duct-derived neogenesis represents an alternative potential source of new β-cells (36; 37), yet both pathways are expected to depend on intact Gα_s_–cAMP signaling. In PGsKO mice, this pathway is inactivated across all epithelial compartments, thereby disabling both routes. Future studies dissecting how Gα_s_ signaling regulates ductal neogenic competence, α-to-β conversion, and β-cell replication could provide valuable therapeutic insight.

In conclusion, pancreatic Gα_s_ is a non-redundant signaling hub whose integrity is essential for postnatal architecture, function, and regenerative capacity. Its deletion leads to a dual endocrine-exocrine failure: disrupted GPCR signal integration in islets and YAP reactivation associated with loss of exocrine polarity. This endocrine-exocrine collapse explains the severe hyperglycemia and growth retardation characteristic of the PGsKO phenotype. Importantly, the unmasking of a latent regenerative response—likely triggered as an adaptive response to chronic hyperglycemia—though insufficient to reverse diabetes in this severe model, opens a promising avenue for therapeutic exploration. Our data suggest that strategically biasing signaling away from specific Gα_s_-coupled GPCRs could be leveraged to promote β-cell regeneration from other pancreatic cell types, offering a novel strategy for diabetes therapy rooted in understanding the pancreas as an integrated organ system.

## Acknowledgements

The authors would like to thank Dr. Omar Coso (IFIBYNE-UBA-CONICET, Argentina) for facilitating the importation of the transgenic mouse lines. The authors would also like to thank Dr. Mariana Judith Elías and Dr. Lirane Moutinho (IFIBYNE-UBA-CONICET, Argentina) for technical assistance with the mouse work. S.A.R.-S. is a career investigator of the Consejo Nacional de Investigaciones Científicas y Técnicas of Argentina (CONICET). This work was supported by the CONICET/NIH 2015 Bilateral Cooperation Programme (PR-512/2016) to J.S.G. and S.A.R.-S. Research in the S.A.R.-S. laboratory was funded by grants from the Agencia Nacional de Promoción Científica y Tecnológica of Argentina (PICT-2019-000374, PICT-2021-00073). The authors also wish to specially acknowledge the Faculty of Exact and Natural Sciences of the University of Buenos Aires for its critical support in sustaining animal facility operations following the suspension of disbursements for the existing national grant supporting this work. M.R., A.R., S.A.T. and A.C.H. are supported by PhD fellowships from the CONICET. J.I.B. was supported by a postdoctoral fellowship from Agencia Nacional de Promoción Científica y Tecnológica of Argentina, and by a travelling fellowship (JCSTF2208815) from The Company of Biologists. During the preparation of this work, the authors used AI-assisted tools (DeepSeek and ChatGPT) to support bioinformatics code adaptation for data visualization and for manuscript grammar checks. Following the use of these tools, the authors formally reviewed the content for accuracy and edited it as necessary. The authors take full responsibility for all content of this publication.

## Author Contributions

J.S.G and S.A.R.-S. conceptualized the work and designed the experiments. M.R., J.I.B., D.J.R., A.R., S.A.T., and V.B. performed wet lab experiments. M.R. and J.I.B. performed quantifications and discussed results with S.A.R-S. M.R., M.Z., and A.C.H. performed bioinformatics analyses. M.R. and S.A.R-S. designed figures and wrote the manuscript, with contributions from all authors. All authors discussed the results, read and approved the final manuscript version. S.A.R-S. is the guarantor of this work and, as such, had full access to all the data in the study and takes responsibility for the integrity of the data and the accuracy of the data analysis.

## SUPPLEMENTARY MATERIAL

## Supplementary FIGURE LEGENDS

**Supplementary Figure 1.**
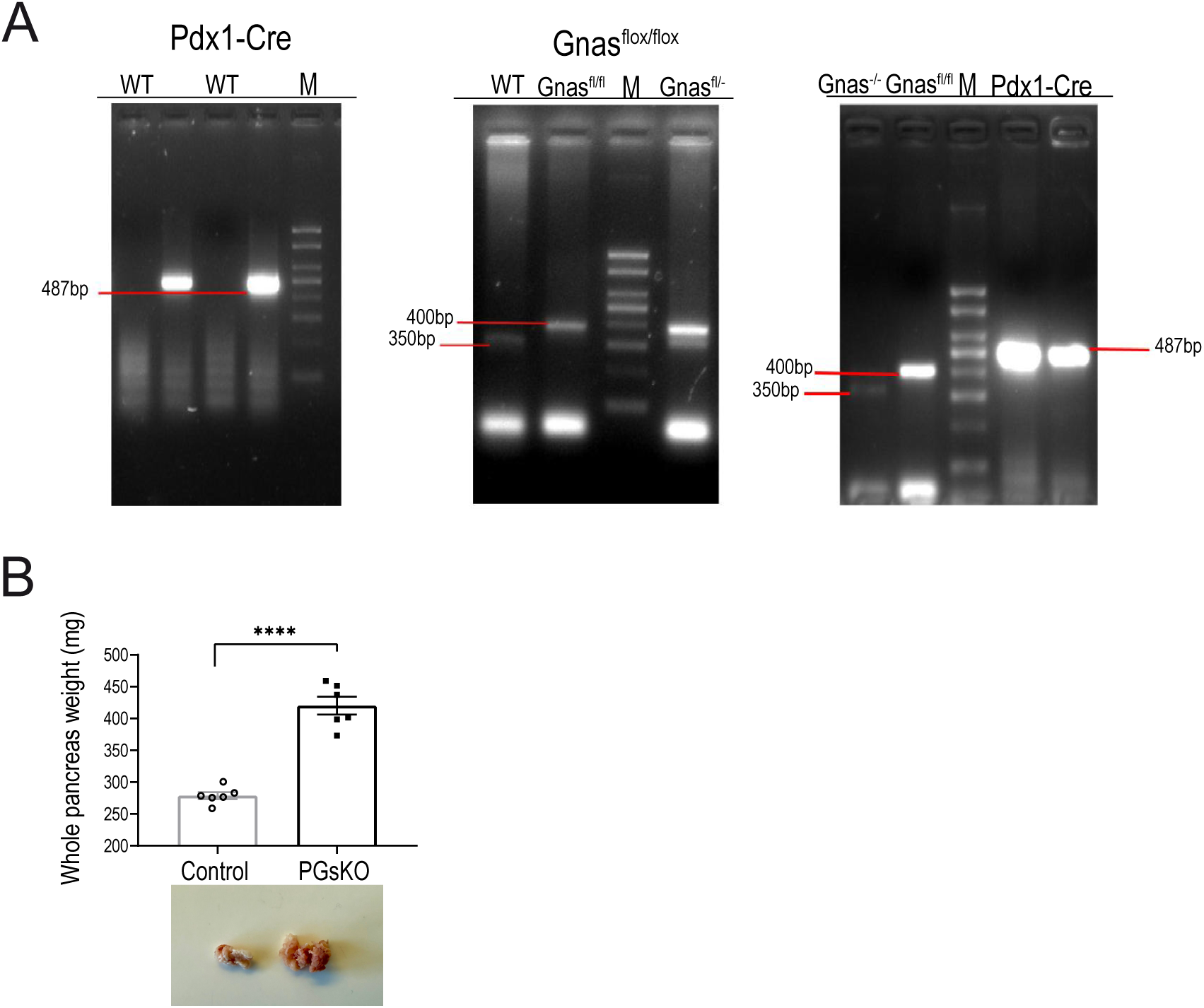
***A:*** Electrophoresis diagram of genomic DNA obtained from tail of Gnas^flox/flox^, Pdx1-Cre^+^ and Gnas^flox/flox^ : Pdx1-Cre^+^ mice by PCR amplification. ***B:*** Quantification of total pancreatic mass in 10-week-old control and PGsKO mice (n=6 mice/group) with a representative photograph of a pancreas for each genotype shown below. Data are expressed as mean ± SEM and statistical analysis was conducted by two-tailed Welch’s t-tests. ****p < 0.0001.

**Supplementary Figure 2.**
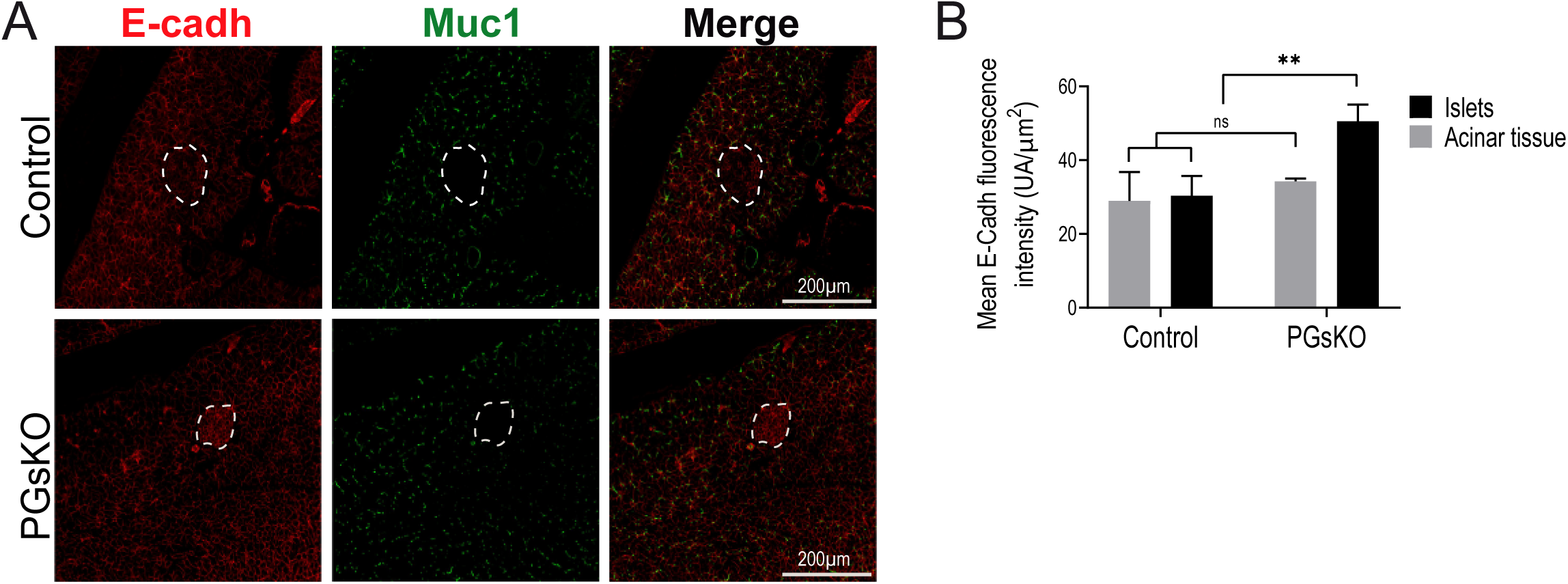
***A:*** Fluorescent labeling of E-cadherin and mucin-1 in pancreatic tissue from 10-week-old PGsKO and control mice. White dashed lines outline the islets. ***B:*** Compartment-specific quantification of mean E-cadherin fluorescence intensity in islets and acinar tissue from 10-week-old control and PGsKO mice (n = 8 control; n = 9 PGsKO). **p < 0.01, ns= not significant.

**Supplementary Figure 3.**
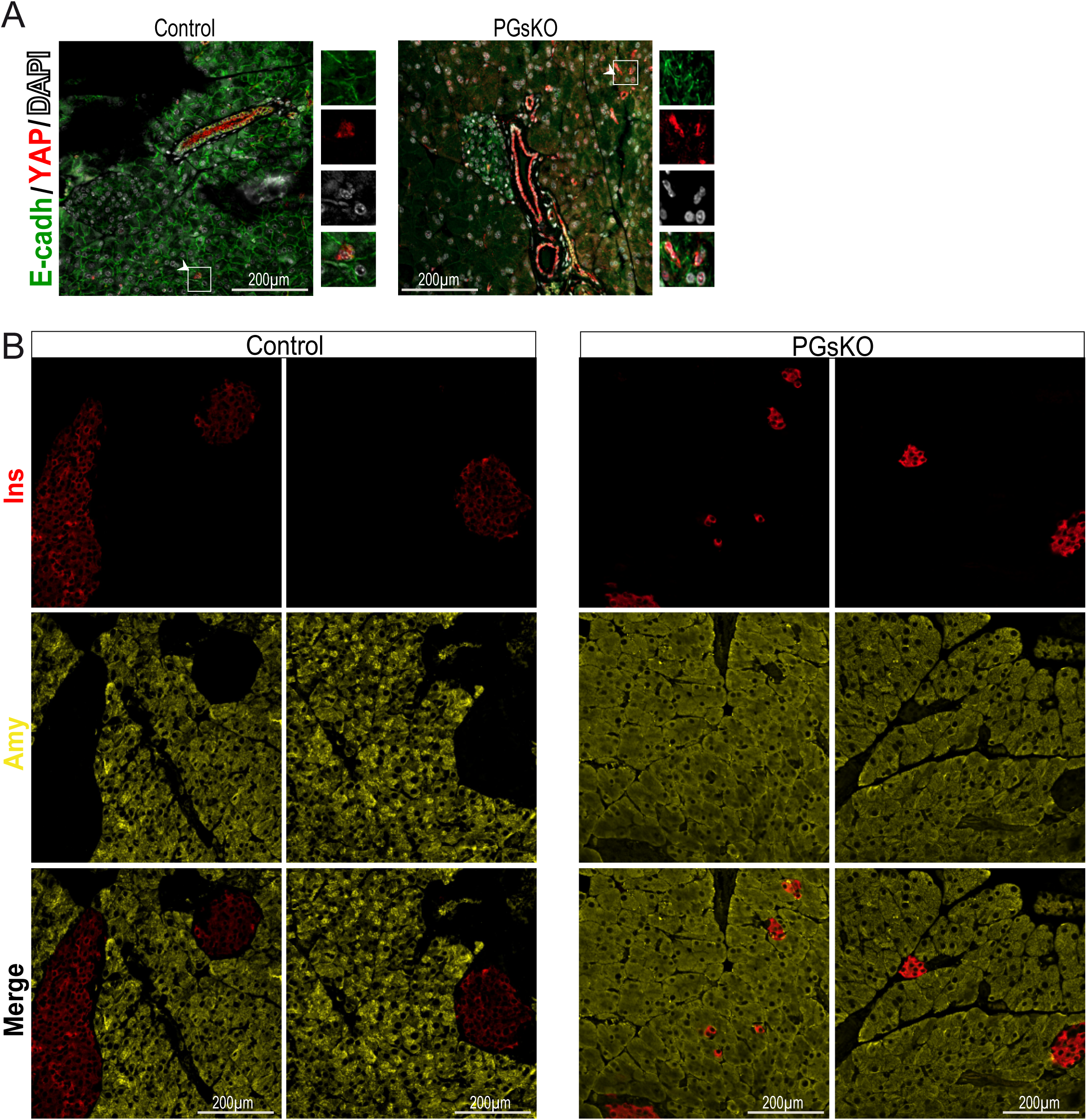
***A:*** Fluorescent staining of pancreatic sections from PGsKO and control mice (10-week-old), labeling E-cadherin, YAP and DAPI; white arrowheads and insets highlight nuclear YAP expression. ***B:*** Representative images of insulin and amylase immunostaining of pancreatic slides of 10-week-old PGsKO and control mice.

**Supplementary Figure 4.**
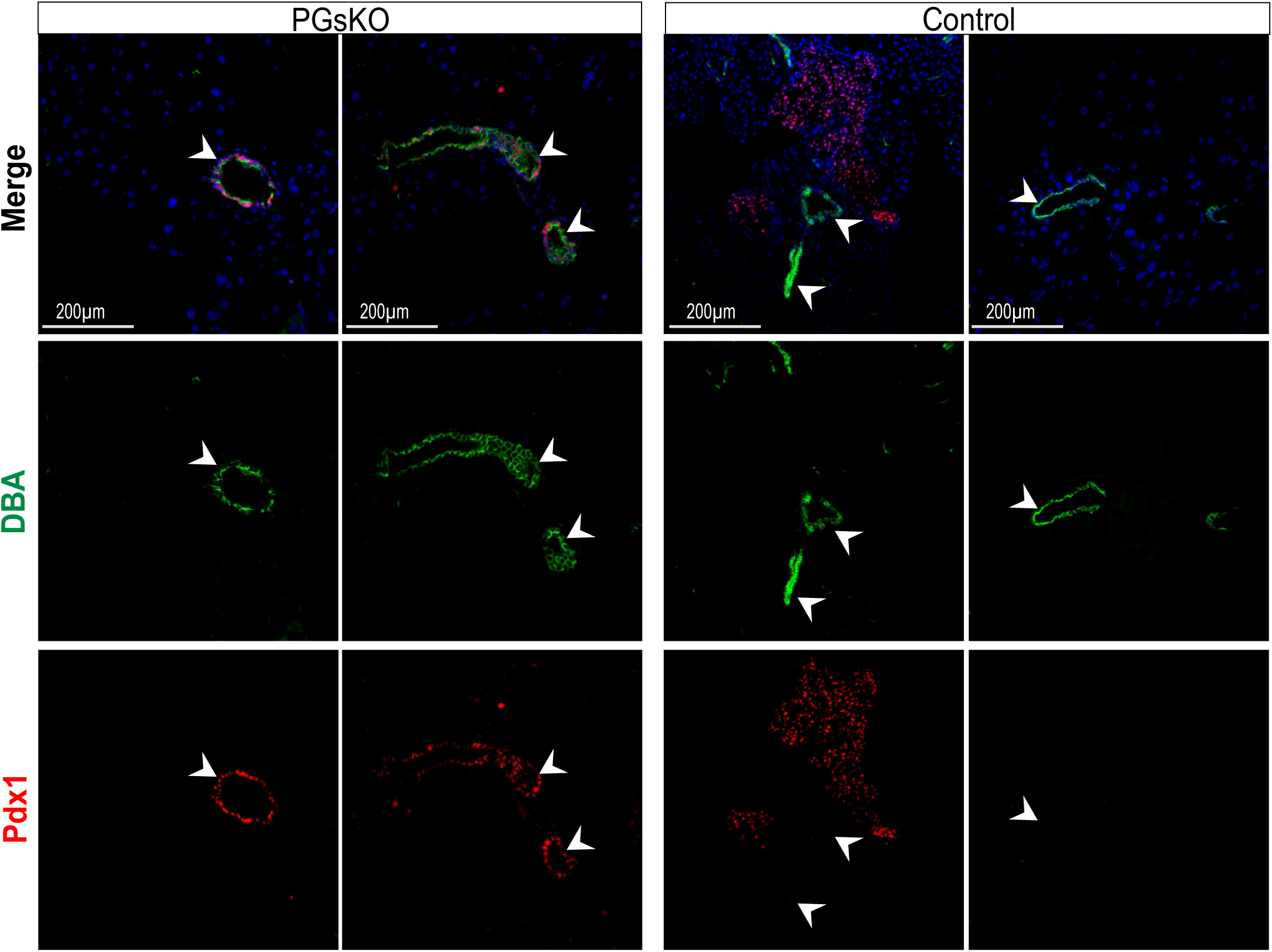
Fluorescent staining of pancreatic sections from PGsKO and control mice (10-week-old), labeling Pdx1, DBA and DAPI with white arrowheads highlighting ducts.

## REFERENCES

1. Skyler JS, Bakris GL, Bonifacio E, Darsow T, Eckel RH, Groop L, Groop P-H, Handelsman Y, Insel RA, Mathieu C, McElvaine AT, Palmer JP, Pugliese A, Schatz DA, Sosenko JM, Wilding JPH, Ratner RE: Differentiation of Diabetes by Pathophysiology, Natural History, and Prognosis. Diabetes 66:241–255, 2017

2. Hogrebe NJ, Ishahak M, Millman JR: Developments in stem cell-derived islet replacement therapy for treating type 1 diabetes. Cell Stem Cell 30:530–548, 2023

3. Oropeza D, Herrera PL: Glucagon-producing α-cell transcriptional identity and reprogramming towards insulin production. Trends in Cell Biology 34:180–197, 2024

4. Bourgeois S, Coenen S, Degroote L, Willems L, Van Mulders A, Pierreux J, Heremans Y, De Leu N, Staels W: Harnessing beta cell regeneration biology for diabetes therapy. Trends in Endocrinology & Metabolism 35:951–966, 2024

5. Drucker DJ: Mechanisms of Action and Therapeutic Application of Glucagon-like Peptide-1. Cell Metabolism 27:740–756, 2018

6. Schreier PCF, Beyerle P, Boulassel S, Beck A, Novikoff A, Reinach PS, Boekhoff I, Breit A, Neuberger A, Müller TD, Serrano AC, Gudermann T, Khajavi N: GLP-1/GIP/GCG receptor triagonist (IUB447) enhances insulin secretion via GLP-1 receptor and Gαq signalling pathway in mice. Diabetologia, 2025

7. Oduori OS, Murao N, Shimomura K, Takahashi H, Zhang Q, Dou H, Sakai S, Minami K, Chanclon B, Guida C, Kothegala L, Tolö J, Maejima Y, Yokoi N, Minami Y, Miki T, Rorsman P, Seino S: Gs/Gq signaling switch in β cells defines incretin effectiveness in diabetes. The Journal of Clinical Investigation 130:6639–6655, 2020

8. Lee Y-S, Lee C, Choung J-S, Jung H-S, Jun H-S: Glucagon-Like Peptide 1 Increases β-Cell Regeneration by Promoting α- to β-Cell Transdifferentiation. Diabetes 67:2601–2614, 2018

9. Wei T, Cui X, Jiang Y, Wang K, Wang D, Li F, Lin X, Gu L, Yang K, Yang J, Hong T, Wei R: Glucagon Acting at the GLP-1 Receptor Contributes to β-Cell Regeneration Induced by Glucagon Receptor Antagonism in Diabetic Mice. Diabetes 72:599–610, 2023

10. Overton DL, Mastracci TL: Exocrine-Endocrine Crosstalk: The Influence of Pancreatic Cellular Communications on Organ Growth, Function and Disease. Frontiers in endocrinology 13:904004, 2022

11. Serra-Navarro B, Fernandez-Ruiz R, García-Alamán A, Pradas-Juni M, Fernandez-Rebollo E, Esteban Y, Mir-Coll J, Mathieu J, Dalle S, Hahn M, Ahlgren U, Weinstein LS, Vidal J, Gomis R, Gasa R: Gsα-dependent signaling is required for postnatal establishment of a functional β-cell mass. Molecular Metabolism 53:101264, 2021

12. Xie T, Chen M, Zhang Q-H, Ma Z, Weinstein LS: β cell-specific deficiency of the stimulatory G protein α-subunit Gsα leads to reduced β cell mass and insulin-deficient diabetes. Proceedings of the National Academy of Sciences 104:19601, 2007

13. Xie T, Chen M, Weinstein LS: Pancreas-specific Gsα deficiency has divergent effects on pancreatic α and β cell proliferation. The Journal of endocrinology 206:261–269, 2010

14. Chen M, Gavrilova O, Zhao W-Q, Nguyen A, Lorenzo J, Shen L, Nackers L, Pack S, Jou W, Weinstein LS: Increased glucose tolerance and reduced adiposity in the absence of fasting hypoglycemia in mice with liver-specific Gsα deficiency. The Journal of Clinical Investigation 115:3217–3227, 2005

15. Cebola I, Rodriguez-Segui SA, Cho CH, Bessa J, Rovira M, Luengo M, Chhatriwala M, Berry A, Ponsa-Cobas J, Maestro MA, Jennings RE, Pasquali L, Moran I, Castro N, Hanley NA, Gomez-Skarmeta JL, Vallier L, Ferrer J: TEAD and YAP regulate the enhancer network of human embryonic pancreatic progenitors. Nat Cell Biol 17:615–626, 2015

16. Heidenreich AC, Bacigalupo L, Rossotti M, Rodríguez-Seguí SA: Identification of mouse and human embryonic pancreatic cells with adult Procr(+) progenitor transcriptomic and epigenomic characteristics. Front Endocrinol (Lausanne) 16:1543960, 2025

17. Pertea M, Kim D, Pertea GM, Leek JT, Salzberg SL: Transcript-level expression analysis of RNA-seq experiments with HISAT, StringTie and Ballgown. Nature Protocols 11:1650–1667, 2016

18. Langmead B, Salzberg SL: Fast gapped-read alignment with Bowtie 2. Nat Methods 9:357–359, 2012

19. Li H, Handsaker B, Wysoker A, Fennell T, Ruan J, Homer N, Marth G, Abecasis G, Durbin R: The Sequence Alignment/Map format and SAMtools. Bioinformatics 25:2078–2079, 2009

20. Robinson JT, Thorvaldsdóttir H, Winckler W, Guttman M, Lander ES, Getz G, Mesirov JP: Integrative genomics viewer. Nature Biotechnology 29:24–26, 2011

21. Ramírez F, Ryan DP, Grüning B, Bhardwaj V, Kilpert F, Richter AS, Heyne S, Dündar F, Manke T: deepTools2: a next generation web server for deep-sequencing data analysis. Nucleic Acids Research 44:W160–W165, 2016

22. Pándy-Szekeres G, Munk C, Tsonkov TM, Mordalski S, Harpsøe K, Hauser AS, Bojarski AJ, Gloriam DE: GPCRdb in 2018: adding GPCR structure models and ligands. Nucleic Acids Research 46:D440–D446, 2017

23. Lin EE, Scott-Solomon E, Kuruvilla R: Peripheral Innervation in the Regulation of Glucose Homeostasis. Trends in Neurosciences 44:189–202, 2021

24. Andersson O, Adams Bruce A, Yoo D, Ellis Gregory C, Gut P, Anderson Ryan M, German Michael S, Stainier DYR: Adenosine Signaling Promotes Regeneration of Pancreatic β Cells In Vivo. Cell Metabolism 15:885–894, 2012

25. Liu L, El K, Dattaroy D, Barella LF, Cui Y, Gray SM, Guedikian C, Chen M, Weinstein LS, Knuth E, Jin E, Merrins MJ, Roman J, Kaestner KH, Doliba N, Campbell JE, Wess J: Intra-islet α-cell Gs signaling promotes glucagon release. Nature Communications 15:5129, 2024

26. Iglesias-Bartolome R, Torres D, Marone R, Feng X, Martin D, Simaan M, Chen M, Weinstein LS, Taylor SS, Molinolo AA, Gutkind JS: Inactivation of a Galphas-PKA tumour suppressor pathway in skin stem cells initiates basal-cell carcinogenesis. Nat Cell Biol 17:793–803, 2015

27. George NM, Day CE, Boerner BP, Johnson RL, Sarvetnick NE: Hippo signaling regulates pancreas development through inactivation of Yap. Molecular and cellular biology 32:5116-5128, 2012

28. Ideno N, Yamaguchi H, Ghosh B, Gupta S, Okumura T, Steffen DJ, Fisher CG, Wood LD, Singhi AD, Nakamura M, Gutkind JS, Maitra A: GNAS(R201C) Induces Pancreatic Cystic Neoplasms in Mice That Express Activated KRAS by Inhibiting YAP1 Signaling. Gastroenterology 155:1593–1607 e1512, 2018

29. Serrill JD, Sander M, Shih HP: Pancreatic Exocrine Tissue Architecture and Integrity are Maintained by E-cadherin During Postnatal Development. Scientific Reports 8:13451, 2018

30. Tixi W, Maldonado M, Chang Y-T, Chiu A, Yeung W, Parveen N, Nelson MS, Hart R, Wang S, Hsu WJ, Fueger P, Kopp JL, Huising MO, Dhawan S, Shih HP: Coordination between ECM and cell-cell adhesion regulates the development of islet aggregation, architecture, and functional maturation. Elife 12:e90006, 2023

31. Gromada J, Chabosseau P, Rutter GA: The α-cell in diabetes mellitus. Nature Reviews Endocrinology 14:694–704, 2018

32. Chandra R, Liddle RA: Cholecystokinin. Current Opinion in Endocrinology, Diabetes and Obesity 14, 2007

33. Antonioli L, Blandizzi C, Csóka B, Pacher P, Haskó G: Adenosine signalling in diabetes mellitus pathophysiology and therapeutic considerations. Nature Reviews Endocrinology 11:228–241, 2015

34. Carboneau BA, Breyer RM, Gannon M: Regulation of pancreatic β-cell function and mass dynamics by prostaglandin signaling. Journal of Cell Communication and Signaling 11:105–116, 2017

35. Courty E, Besseiche A, Do TTH, Liboz A, Aguid FM, Quilichini E, Buscato M, Gourdy P, Gautier JF, Riveline JP, Haumaitre C, Buyse M, Fève B, Guillemain G, Blondeau B: Adaptive β-Cell Neogenesis in the Adult Mouse in Response to Glucocorticoid-Induced Insulin Resistance. Diabetes 68:95–108, 2019

36. Fernández Á, Casamitjana J, Holguín-Horcajo A, Coolens K, Mularoni L, Guo L, Hartwig O, Düking T, Vidal N, Strickland LN, Pasquali L, Bailey-Lundberg JM, Rooman I, Wang YJ, Rovira M: A Single-Cell Atlas of the Murine Pancreatic Ductal Tree Identifies Novel Cell Populations With Potential Implications in Pancreas Regeneration and Exocrine Pathogenesis. Gastroenterology 167:944–960.e915, 2024

37. Doke M, Álvarez-Cubela S, Klein D, Altilio I, Schulz J, Mateus Gonçalves L, Almaça J, Fraker CA, Pugliese A, Ricordi C, Qadir MMF, Pastori RL, Domínguez-Bendala J: Dynamic scRNA-seq of live human pancreatic slices reveals functional endocrine cell neogenesis through an intermediate ducto-acinar stage. Cell Metabolism 35:1944–1960.e1947, 2023

